# Common garden experiments suggest terpene-mediated interactions between phyllosphere microbes and *Cryptomeria japonica*

**DOI:** 10.1101/2025.05.04.652152

**Authors:** Satoyoshi Ishizaki, Tetsuo I. Kohyama, Yuki Ota, Takuya Saito, Yoshihisa Suyama, Yoshihiko Tsumura, Tsutom Hiura

**Affiliations:** Department of Ecosystem Studies, Graduate School of Agricultural and Life Sciences, The University of Tokyo, Tokyo 113-8657, Japan; National Institute for Environmental Studies, Tsukuba 305-8506, Japan; Field Science Center, Graduate School of Agricultural Science, Tohoku University, Osaki 989-6711, Japan; Institute of Life and Environmental Sciences, University of Tsukuba, Tsukuba 305-8572, Japan

## Abstract

Plant–microbe interactions in the phyllosphere provide invaluable information on plant ecology, with implications for ecosystem functioning and plant–atmosphere feedbacks. The composition of phyllosphere microbial communities varies significantly depending on host lineages, geographic regions, and climatic conditions. However, the factors driving these variations in interactions with plants remain poorly understood. Biogenic volatile organic compounds (BVOCs) emitted by plants may be important in these interactions. Here, we quantified the composition of phyllosphere microbial communities and terpene emissions from leaves of Japanese cedar (*Cryptomeria japonica*) trees grown in two common gardens from cuttings collected from natural populations across Japan. Amplicon sequencing revealed that bacterial and fungal communities differed significantly between gardens and among host population origins. Analysis of BVOC profiles showed that the camphene emission rate was associated with bacterial community composition, whereas that of *β*-farnesene was linked to fungal community composition. The relative abundances of certain putative plant pathogens and the emission rates of most monoterpenes were correlated with the climatic conditions at the origin sites of cedar trees. These findings highlight the intricate relationships between phyllosphere microbial communities and terpene emission from host trees and suggest the role of climatic factors in shaping these interactions.

## Introduction

The aerial surface of plant leaves, called the phyllosphere, harbors a diverse array of microbes including bacteria and fungi^1^. Interactions between phyllosphere microbes and host plants have a considerable impact on plant physiology and ecology. Phyllosphere microbes can influence plant fitness by causing or suppressing disease^2–5^ or by promoting plant growth and stress tolerance^6,7^. Plant defense against pathogens and the counteradaptation by the latter further complicate plant–microbe interactions^7–10^. Studies have even suggested that these interactions contribute to ecosystem functioning, including nutrient cycling^11–15^ and ecosystem productivity^12,15,16^. Microbial infection may also affect plant–atmosphere feedbacks through microbial metabolization of gases emitted by plants^14,17–19^. Studies of plant–microbe interactions can lead to insights into key issues in ecology, such as the evolution of plant traits, drivers of ecosystem functions, and biosphere–atmosphere interactions.

Shifts in the composition of the phyllosphere microbial community are significant indicators of changes in plant–microbe interactions^5,15,20^. This composition is affected by environmental factors including climate^21,22^, season^21,23^, land use^24,25^, location^21,26^, and geographic distance^24^ as well as by host-related factors such as species^21,27,28^ and traits^21,29^. The effect of host genotypes on microbial composition in the phyllosphere has been investigated using known characterized mutants or accessions^23,26,30^. However, further investigation of genetic and phenotypic variation within wild plant species is necessary to elucidate the interactions and coevolution of phyllosphere microbes with plants.

Biogenic volatile organic compounds (BVOCs) are plant-derived secondary metabolites that affect plant–microbe interactions^10,31^. BVOCs affect plant resistance to microbes by directly inhibiting microbial growth by damaging microbial cell membranes^32,33^ and inducing plant immune responses^34^. On the other hand, some microbes metabolize BVOCs stored in or emitted from plants and use them as a carbon source, suggesting their adaption to plant resistance mechanisms^13,14,35,36^. While many laboratory studies have suggested a relationship between pathogen-induced BVOC emission by plants and phyllosphere microbes^34,37,38^, few have investigated the roles of constitutive BVOC emission by plants under natural conditions in plant–microbe interactions^39,40^. Significant inter- and intraspecific variations in both the composition of the phyllosphere microbial community and emission rates of BVOCs have been reported for wild plants^39,41,42^. Elucidating the drivers and interrelationships of these variations is a challenge in understanding plant–microbe interactions mediated by plant BVOC emission.

BVOC emission can also have a significant impact on atmospheric chemical processes and climate. Oxidation of emitted BVOCs can indirectly affect the radiative balance by increasing the lifetime of certain greenhouse gases due to a decrease in oxidative capacity in the atmosphere^43^. Secondary organic aerosols produced by BVOC oxidation act as cloud condensation nuclei and may affect regional climate^44^. Guenther et al.^45^ estimated that terrestrial plants emit about 1 billion tons of BVOCs per year, of which about 92% is emitted by forest trees. These estimates are based on the model of the standardized rates of BVOC emission by plants. However, this model does not fully account for variations in the amounts and composition of emitted BVOCs among and within plant species or the influences of plant interactions with microbes. Therefore, elucidating the relationship between plant–microbe interactions and plant BVOC emissions is important for understanding the plant–aerosol– climate feedbacks.

Wild populations of Japanese cedar, *Cryptomeria japonica* (Cupressaceae), are widely distributed in Japan, ranging from Yakushima Island, the southern limit of their distribution, to the northern limit of Ajigasawa (Aomori Prefecture)^46^ (Supplementary Fig. S1 and Table S1). As one of the major silvicultural tree species in Japan, *C. japonica* accounts for approximately 20% of the total forest area, including plantations^47^.

Japanese cedar emits substantial amounts of terpenes, including monoterpenes (MTs), sesquiterpenes (SQTs), and diterpenes (DTs)^48–50^. In comparison with MTs, SQTs and DTs have high molecular weights and are highly reactive in the atmosphere^51,52^. Considering the large numbers of *C. japonica* in Japan’s forests, these terpenes may have a non-negligible impact on the surrounding atmospheric environment. A study based on a common garden experiment reported significant intraspecific variation in the composition of terpenes stored in and emitted from the leaves of Japanese cedars^50^. Intraspecific genetic variation in Japanese cedar has been well studied, revealing a geographic genetic structure with four main clades: the Pacific Ocean side clade, two Sea of Japan side clades, and the Yakushima Island clade^53–55^.

Intraspecific variation in physiological and morphological traits of Japanese cedar has been revealed by common garden experiments, suggesting adaptation to the original environments^56–58^. The reasons for differences in the quantity and composition of stored and emitted terpenes among Japanese cedar populations are still unclear. However, these differences may be influenced by the local climate and pathogen communities at the original locations of the populations^50^. The problem is that the pathogen community data used in that study were based on previously published information, and no quantitative data were obtained on the actual microbial community on the leaf surface of Japanese cedar. To clarify whether the intraspecific variations in terpene emissions observed in this species reflect the variations in the phyllosphere microbial community, quantitative investigations of both microbial communities and BVOCs emitted by the host are necessary.

Here, we investigated the phyllosphere microbial communities of Japanese cedar by amplicon sequencing using cedar trees that have genetic variations and are grown in two common gardens. We also measured terpene emission rates in some of these trees in one common garden. Our objectives were (1) to clarify the variation in the phyllosphere microbial community composition among Japanese cedar populations, and (2) to elucidate the relationship between terpene emission rates and these microbial compositions. Our study provides new insights into the mechanisms of plant–microbe interactions mediated by host-emitted BVOCs.

## Results

### Composition of the phyllosphere microbial community of Japanese cedar

Amplicon sequencing identified 2933 bacterial amplicon sequence variants (ASVs) and 5040 fungal ASVs across all samples from the phyllosphere microbial communities of Japanese cedar planted in the common gardens. After rarefaction, the richness of bacterial ASVs (9099 reads) was 89 ± 64 (mean ± SD) and that of fungal ASVs (17,427 reads) was 136 ± 121. In the bacterial communities, Proteobacteria (58.9% of all reads), Acidobacteriota (6.8%), Myxococcota (5.9%), Verrucomicrobiota (4.6%), Bacteroidota (3.0%), Firmicutes (2.5%), and Actinobacteriota (1.7%) were most abundant (Fig. 1a, b). The relative abundance of Myxococcota tended to be higher in the Kawatabi common garden, whereas that of Acidobacteriota was higher in the Tsukuba common garden. In the fungal communities, Dothideomycetes (32.4% of all reads), Eurotiomycetes (30.7%), Leotiomycetes (9.2%), Lecanoromycetes (7.6%), Sordariomycetes (2.6%), Cystobasidiomycetes (2.5%), Agaricomycetes (2.1%), and Tremellomycetes (1.8%) were most abundant (Fig. 1c, d). The relative abundance of Leotiomycetes and Lecanoromycetes tended to be higher in the Tsukuba common garden than in the Kawatabi common garden.

**Fig. 1.**
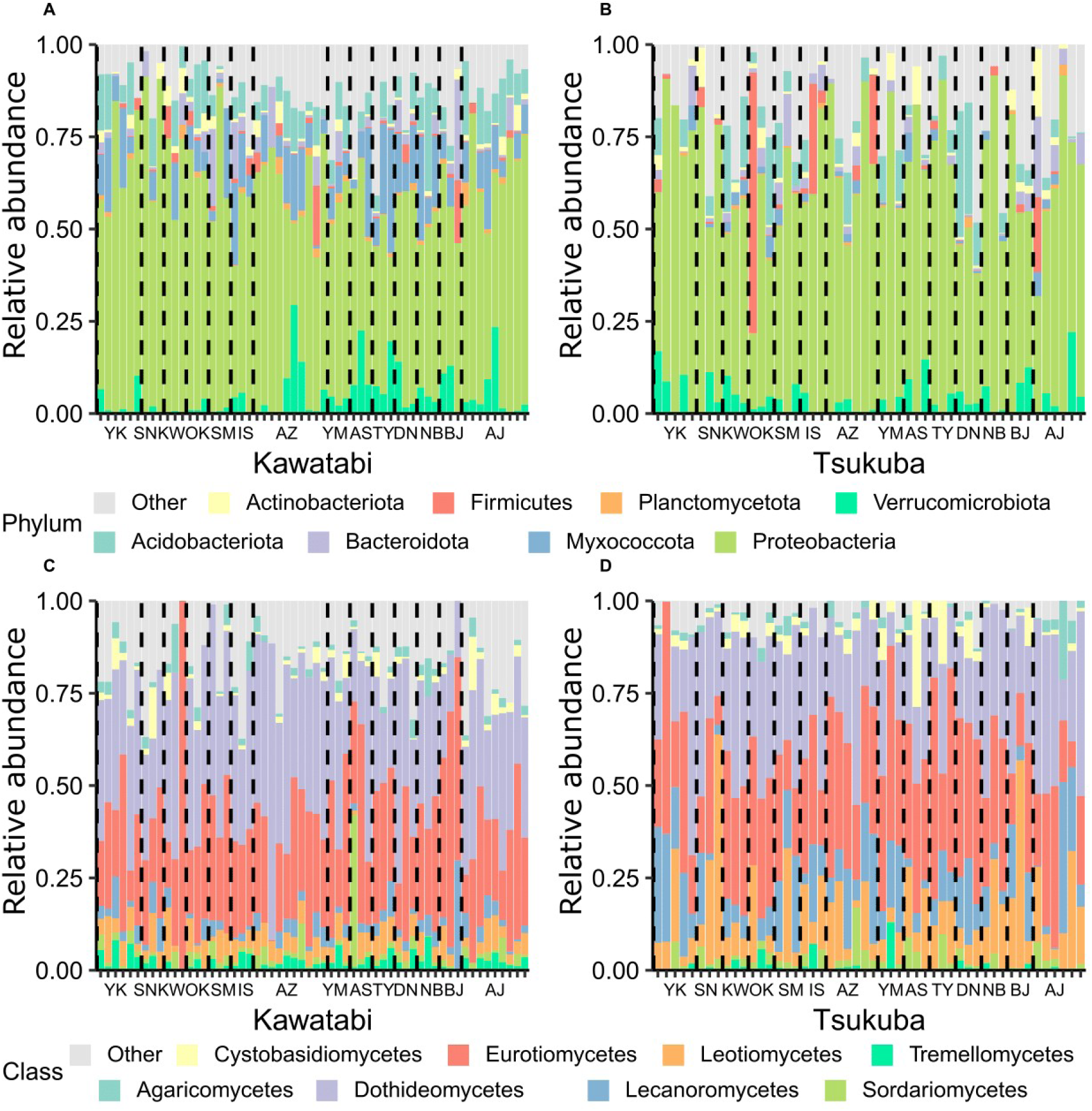
Compositions of the phyllosphere microbial communities of Japanese cedars planted in the common gardens. Each bar represents the phyllosphere microbial community of each individual at (**A, C**) Kawatabi and (**B, D**) Tsukuba. Bar arrangement is based on the value of the climatic PC1 (from high to low) at the origin site of each individual. (**A, B**) Relative abundances of eight most abundant bacterial phyla. (**C, D**) Relative abundances of eight most abundant fungal classes. The original population of each sample is indicated under the bar (see Supplementary Fig. S1 and Table S1 for the names and locations).

The composition of the bacterial and fungal communities was significantly influenced by the locations of the common gardens and host populations (Fig. 2). In the bacterial community, 38.3% of the total variation was explained by the common garden (PERMANOVA: *R*^2^ = 0.130, *p* = 0.001), host population (*R*^2^ = 0.124, *p* = 0.014), and their interaction (*R*^2^ = 0.128, *p* = 0.009). In the fungal community, 40.1% of the total variation was explained by the common garden (PERMANOVA: *R*^2^ = 0.176, *p* = 0.001), host population (*R*^2^ = 0.116, *p* = 0.028), and their interaction (*R*^2^ = 0.115, *p* = 0.033).

**Fig. 2.**
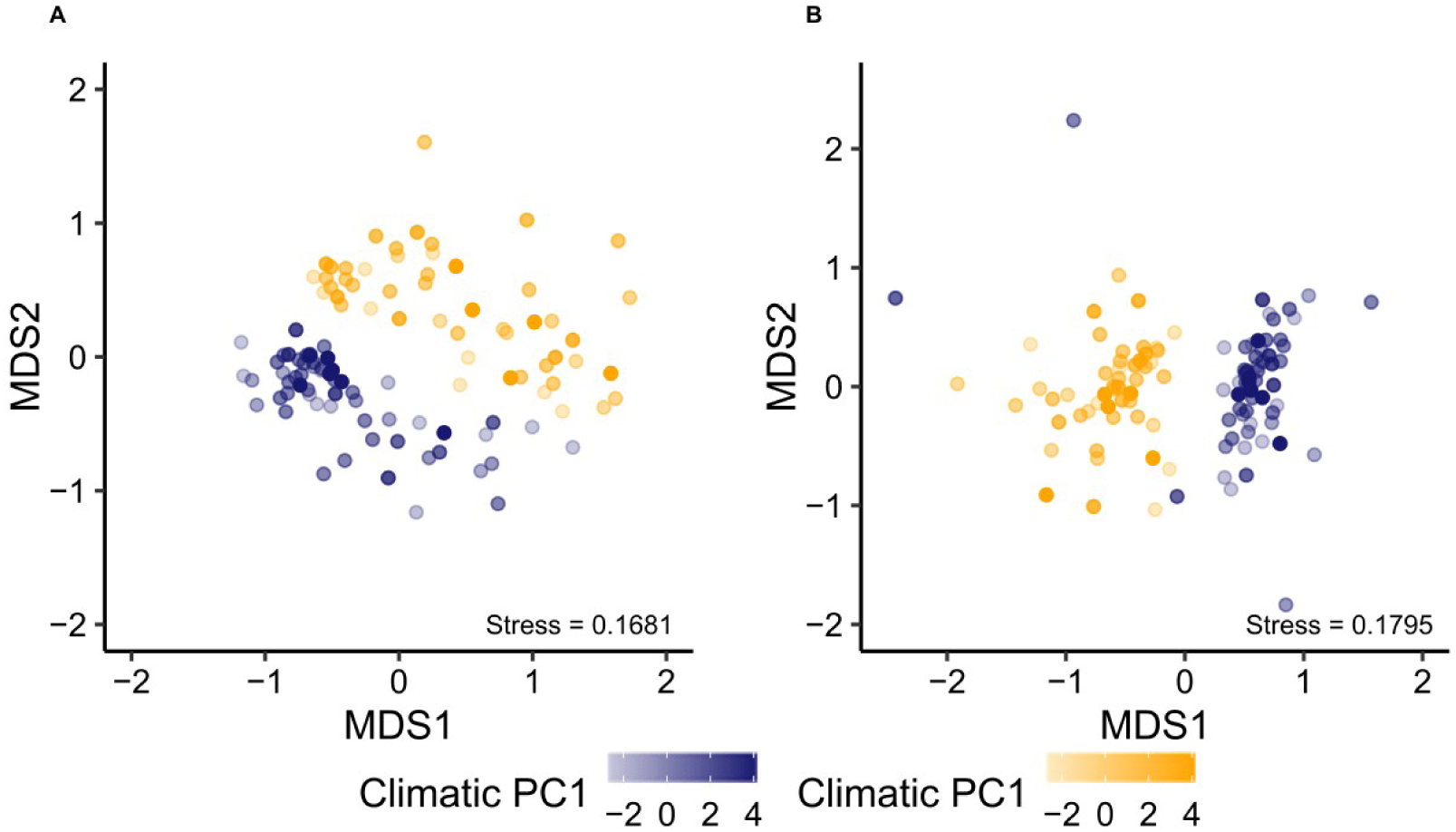
Differences in the composition of the phyllosphere (**A**) bacterial and (**B**) fungal communities of Japanese cedar between the common gardens at Kawatabi (blue) and Tsukuba (orange). Differences were visualized using nonmetric multidimensional scaling (NMDS) based on Bray–Curtis dissimilarities. The density of color represents the climatic PC1 at the origin site of each tree. MDS1 indicates the first ordination axis generated by NMDS and MDS2 indicates the second.

To investigate how climatic conditions at the original locations of the host populations influence their effects on the composition of the microbial communities, we obtained the climatic data^59–62^ at these locations. A principal component analysis (PCA) of these data revealed that 73.8% of the total variation was explained by the first two PCs (Supplementary Fig. S2), which were used for the subsequent analysis. Climatic PC1, which explained 50.57% of the total variation, corresponded to higher annual mean temperature and mean monthly potential evaporation and corresponded to lower standard deviation of the monthly mean temperatures and the sum of the monthly precipitation for the months with mean temperatures below 0 °C, suggesting that higher values of PC1 correspond to a warmer and less snowy climate. Climatic PC2, which explained 23.19% of the total variation, corresponded to higher mean monthly near-surface relative humidity and annual precipitation and a lower coefficient of variation of the monthly precipitation estimates, suggesting that higher values of PC2 correspond to higher amounts of precipitation and its stability.

The total explained variation in microbial community composition was reduced when climatic PC1 and PC2 at the host origin sites were used as explanatory variables instead of the host population. The result of PERMANOVA testing the effects of climatic PCs and their interactions with the locations of the common gardens on the variation in bacterial community composition was as follows: *R*^2^ = 0.014 and *p* = 0.044 for PC1, *R*^2^ = 0.011 and *p* = 0.136 for the interaction between PC1 and common garden, *R*^2^ = 0.009 and *p* = 0.335 for PC2, *R*^2^ = 0.010 and *p* = 0.160 for the interaction between PC2 and common garden. For the total variation in fungal community composition, the result of PERMANOVA was *R*^2^ = 0.010 and *p* = 0.126 for PC1, *R*^2^ = 0.007 and *p* = 0.671 for the interaction between PC1 and common garden, *R*^2^ = 0.009 and *p* = 0.277 for PC2, and *R*^2^ = 0.008 and *p* = 0.346 for the interaction between PC2 and common garden. Of the detected fungal genera, 11 have been described as Japanese cedar pathogens^63^ (Supplementary Table S2). The aggregated abundances of these putative pathogenic fungi after rarefaction were negatively correlated with climatic PC1 at the host origin sites (Wald test: *β* = −0.423 ± 0.158, *z* = −2.676, *p* = 0.007; Fig. 3); in particular, negative correlation was found for the abundances of white rot fungi. However, the influence of common garden and climatic PC2 on these abundances was not statistically significant (both *p* > 0.1).

**Fig. 3.**
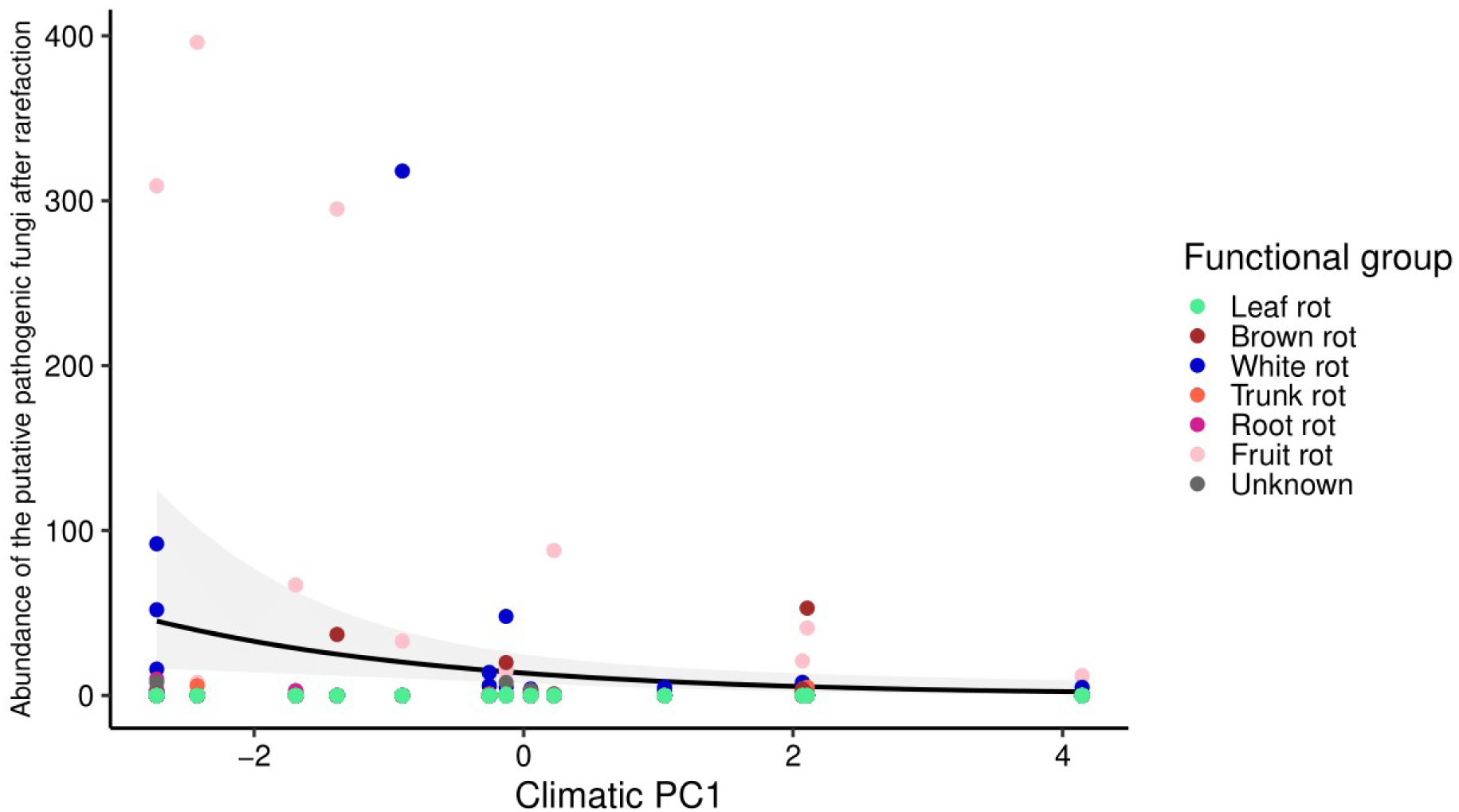
Relationship between the abundance of the putative pathogenic fungi after rarefaction and climatic PC1 at the origin sites of the host trees. The abundances of the fungi are displayed. They were aggregated on the basis of their putative functional groups (color of the points). The solid line represents fitted values from a negative binomial generalized linear model, where the aggregated abundance of all guilds was modeled as a function of climatic PC1. Shaded areas indicate the 95% confidence intervals of the model predictions.

### Relationships between terpene emissions, phyllosphere microbial community composition, and climate

Japanese cedar trees in the Kawatabi common garden emitted 12 MTs, one SQT, and one DT (Fig. 4a). The DT was excluded from the subsequent analysis since only one tree emitted it. PCA showed that PC1 and PC2 explained 70.8% of the total variation in the log-transformed basal emission rates of terpenes (Fig. 4b). Terpene PC1 corresponded to higher basal emission rates of most MTs and was significantly positively correlated with climatic PC1 and PC2 at the tree origin sites (Student’s *t*-test: climatic PC1, *β* = 0.459 ± 0.189, *t* = 2.436, *p* = 0.023; climatic PC2, *β* = 0.874 ± 0.417, *t* = 2.094, *p* = 0.047; Supplementary Fig. S3). Terpene PC2 corresponded to the higher basal emission rates of some MTs and *β*-farnesene but was not significantly correlated with the climatic PCs (Student’s *t*-test: climatic PC1, *β* = −0.009 ± 0.108, *t* = −0.084, *p* = 0.934; climatic PC2, *β* = −0.185 ± 0.230, *t* = −0.803, *p* = 0.430; Supplementary Fig. S3).

**Fig. 4.**
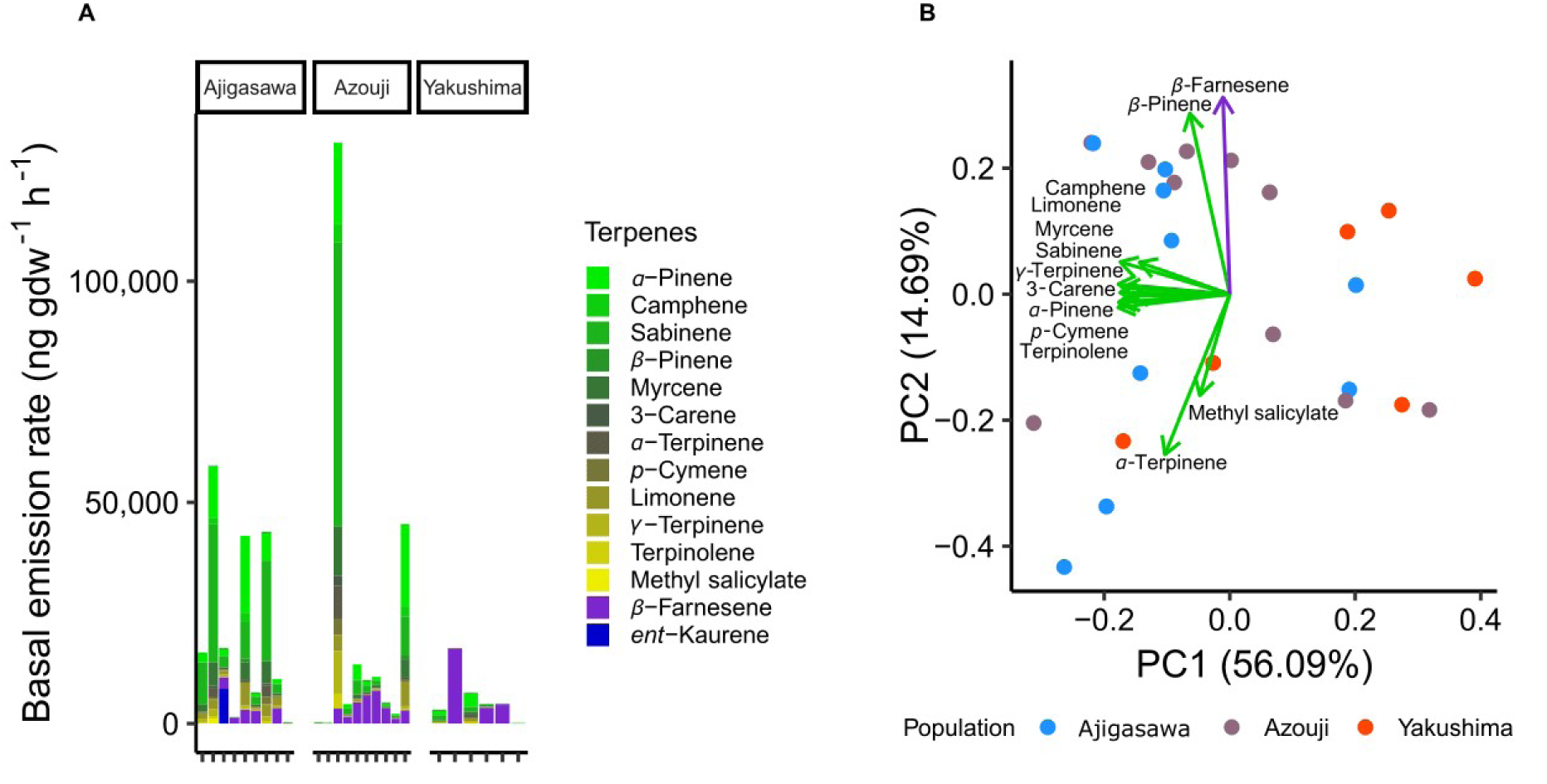
Terpene emission profiles of Japanese cedar in the common garden at Kawatabi. (**A**) Basal emission rates of monoterpenes (green to yellow), a sesquiterpene (purple), and a diterpene (blue) emitted from each cedar individual originating from three populations. (**B**) Principal component analysis of log-transformed basal emission rates of terpenes.

Significant correlations were detected between the basal emission rate of camphene and bacterial community composition (*R*^2^ = 0.448, *p* = 0.003; Fig. 5a). The sum of the basal emission rates of MTs was marginally correlated with bacterial composition (*R*^2^ = 0.187, *p* = 0.098). The basal emission rate of *β*-farnesene was significantly correlated with fungal community composition (*R*^2^ = 0.239, *p* = 0.046; Fig. 5b). The higher basal emission rate of camphene and higher sum of the basal emission rates of MTs corresponded to higher abundances of Actinobacteriota, Firmicutes, Bacteroidota, and Cyanobacteria. The higher basal emission rate of *β*-farnesene corresponded to higher abundances of Taphrinomycetes, Tremellomycetes, and Agaricomycetes.

**Fig. 5.**
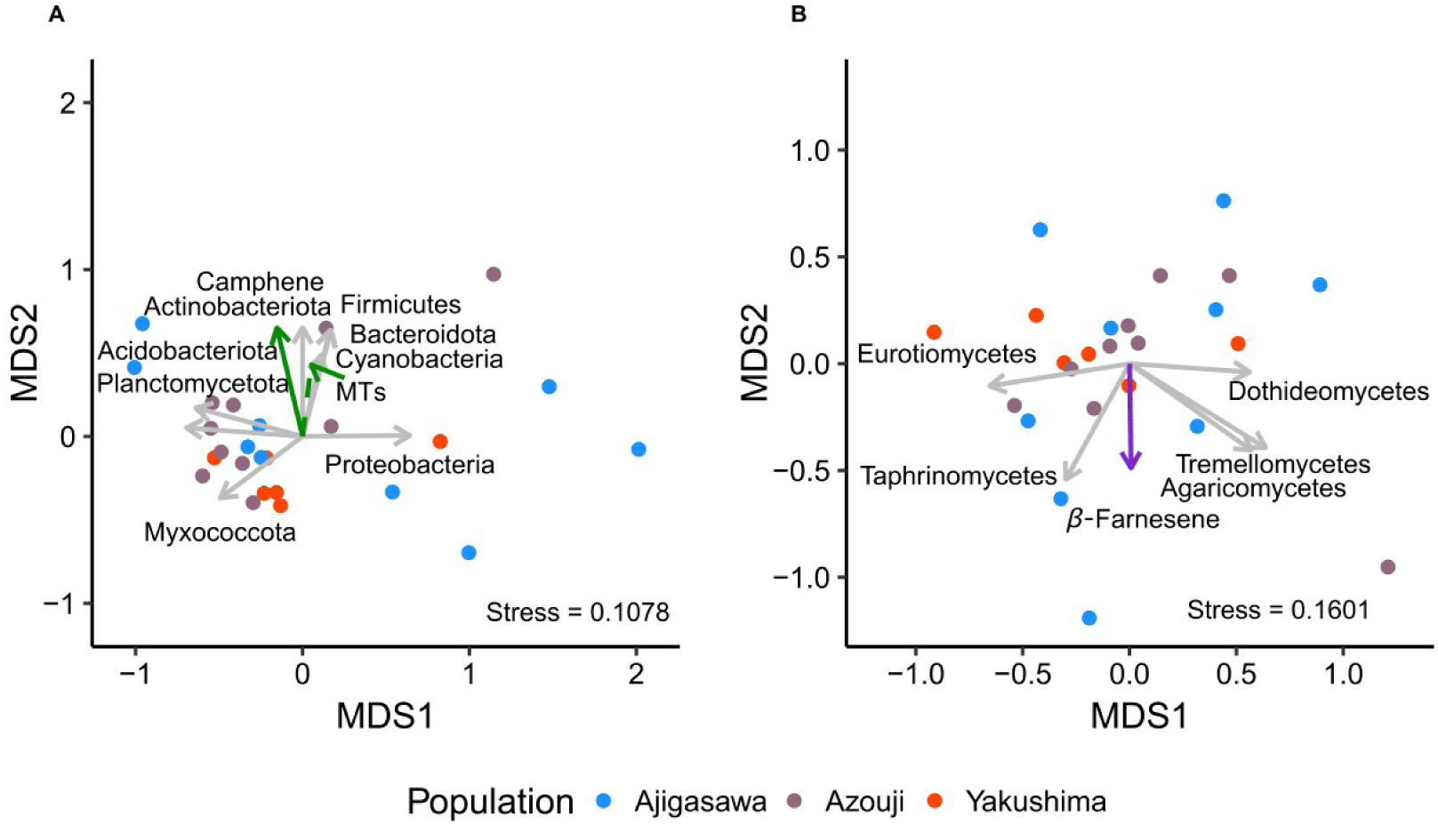
Vector fitting of log-transformed basal emission rates of terpenes onto NMDS coordinates constructed from the compositions of the phyllosphere (**A**) bacterial and (**B**) fungal communities of Japanese cedar from three populations (blue: Ajigasawa, grey: Azouji, orange: Yakushima). (**A**) Bacterial phyla and (**B**) fungal classes whose relative abundances were significantly correlated with the NMDS coordinates are also shown. Only loading vectors of significantly correlated terpenes and microbial taxa are shown (solid: *p* < 0.05, dashed: *p* < 0.1).

## Discussion

Using common garden experiments, we revealed differences in the composition of both bacterial and fungal communities in the phyllosphere among wild populations of Japanese cedar (Fig. 2). The data suggest that genetic variation among these populations influences microbial composition in the phyllosphere. This result is consistent with those of previous studies on grasses^23,26^ and broad-leaved trees^64^. Although climate at the origin sites of cedar populations explained only a relatively small proportion of the total variation in overall microbial compositions, the relative abundances of putative pathogenic fungi were negatively correlated with warm and less snowy climate at the host origin sites (Fig. 3), suggesting the influence of climate on coevolution of some phyllosphere microbes and Japanese cedar. Several studies have suggested a correlation between microbial community composition and the environment, including climate^65–69^. It is possible that changes in distribution among fungi coupled to climatic shifts may have influenced the coevolution of these fungi with interacting plants. The relative abundances of the putative white rot fungal genera (*Trametes*, *Phanerochaete*, *Pholiota*, and *Phellinus*) were negatively correlated with warmth at the host origin sites (Fig. 3). Pathogens of these genera have been reported in wild Japanese cedar populations, mainly in the northern region of Honshu Island^50^, which is consistent with the results of this study. Field-grown Japanese cedar trees from cold-climate populations have been suggested to store higher amounts of terpenes^50^, and these pathogens may be better adapted than other microbes to this defense trait.

Our data suggest that the basal emission rates of some terpenes in Japanese cedar may be related to the composition of the phyllosphere microbial community. The basal emission rate of the MT camphene was correlated with the composition of the bacterial community (Fig. 5a). However, its amount was smaller than those of other terpenes such as *α*-pinene and sabinene. The sum of the basal emission rates of MTs was also marginally correlated with bacterial composition, and the direction of its loading vector was similar to that of camphene. These results suggest the regulation of bacterial communities in the phyllosphere of Japanese cedar by the emission of some MTs, possibly by that of camphene. The emission of multiple terpenes is simultaneously induced by plant defense signaling pathways^10,17,70^. Camphene, together with other MTs *α*-pinene and *β*-pinene, has been suggested to play a pivotal role in the induction of defense responses against bacterial pathogens in *Arabidopsis thaliana*^34^. Given the correlation between the basal emission rate of camphene and those of most of the other MTs (Fig. 4), multiple MTs may collectively control the bacterial communities on Japanese cedar. It is also possible that the detection limits of terpene emission affected our results, since many of the determined emission rates of camphene were low and this compound was not detected in several samples. In addition, the outliers in the camphene emission data may have influenced the correlation test. Furthermore, the small number of populations used for BVOC measurements may have been a limiting factor, and further investigation is needed.

The basal emission rate of the SQT *β*-farnesene was correlated with the composition of the fungal community. The higher basal emission rate of the *β*-farnesene corresponded to the higher relative abundances of Taphrinomycetes, Tremellomycetes, and Agaricomycetes (Fig. 5b). *β*-Farnesene is emitted upon insect herbivory in maize^71^ and pathogen infection in wheat^72^. Japanese cedar may also use *β*-farnesene as a defense substance against fungal infection. Hiura et al.^50^ suggested a correlation between the composition of the fungal pathogen community in each cedar population and the relative basal emission rates of the SQT *α*-farnesene and the DT *ent*-kaurene. Taken together, their data and the results of this study suggest that Japanese cedar may be adapting to fungal biotic stresses by using relatively high-molecular-weight terpenes such as SQTs and DTs.

Some microbes are able to regulate or use plant defenses^10,73^. Firmicutes was one of the bacterial phyla whose relative abundance was higher on the hosts with higher basal emission rate of camphene (Fig. 5a). Some strains of Firmicutes induce immunity in tomato and are relatively abundant in plants with enhanced defense^74^. The induction of plant immunity often leads to the emission of terpenes^17,34,70^. However, the elevated level of camphene emission observed in this study may not be induced by bacteria, as the MTs emitted by Japanese cedar were suggested to be released primarily from storage pools^75^. Nevertheless, the possibility remains that Firmicutes may tolerate cedar defenses mediated by enhanced camphene emission. Taphrinomycetes, whose relative abundance was positively related to the basal emission rate of *β*-farnesene (Fig. 5b), include the genus *Taphrina*. All known *Taphrina* species are plant pathogens^76^ and have various functional genes required for plant infection by these fungi, such as genes encoding enzymes involved in plant hormone biosynthesis or degradation of plant cell walls^77^. The positive correlations between the relative abundances of these microbes and the emissions of some terpenes may indicate microbial adaptation to the phyllosphere environment of Japanese cedar. Further investigation is needed to identify the microbial functions that support the BVOC-mediated interaction with the cedar.

We demonstrated that the locations of common gardens were the primary factor influencing microbial composition in the phyllosphere of Japanese cedar (Fig. 2). Multiple factors may contribute to the observed differences in microbial composition between the gardens. Geographic distance may affect the community assembly of local microbes, particularly those with limited dispersal capabilities^24,68^. Fungi have larger spores^78^ and are more limited in dispersal than bacteria^68,79,80^, which may explain the greater effect of common garden on the fungal composition in this study. The two gardens differed in their local environments since they are located at different latitudes and the surrounding land use is also very different: Kawatabi is surrounded by a forest, whereas Tsukuba is surrounded by farmland and an urban area. These differences in the local environment, such as climate^21,22^ and land use^24,81^, may also have influenced the microbial composition in the gardens.

Although relatively few populations were used, our study suggested a negative correlation between the basal emission rates of most MTs and warm, humid, and less snowy climates at the original locations of the populations. Hiura et al.^50^ suggested that the amounts of stored, but not emitted, MTs in Japanese cedar are negatively correlated with warm and less snowy climates, suggesting the existence of climate clines. Partial pressure calculated from the stored amounts of MTs in Japanese cedar was suggested to be positively correlated with their emission rates from the same leaf^75^. Therefore, the similar pattern of variations among Japanese cedar populations in stored and emitted MTs may be attributed to the emission of MTs from storage pools in leaves. We expect that consideration of both abiotic and biotic environmental conditions will further our understanding of the mechanisms of BVOC emission in plants in natural environments.

In conclusion, our findings revealed differences in the phyllosphere microbial community compositions of *C. japonica* both between the common gardens and among wild populations, as well as the correlation between the compositions and basal emission rates of some terpenes by host trees. The results also suggest that the local climate at the origin sites of the Japanese cedar populations is a significant factor influencing variations in the phyllosphere microbial communities and terpene emission rates in this species. This study is among the few to demonstrate in outdoor settings the relationship between the emission patterns of BVOCs and both abiotic and biotic environments in trees with substantial BVOC emissions. Further investigation is required to elucidate the mechanisms of plant–microbe interactions mediated by BVOC emission and the impact of biotic interactions on plant–atmosphere feedbacks.

## Methods

### Sampling sites and collection of phyllosphere microbiome

We collected leaves at a height of about 1.5 m from cedar trees (about 5–6 m tall) on 2 June 2023 in the common garden at Kawatabi Field Science Center, Tohoku University (38.78°N, 140.73°E) and on 26 June 2023 in the common garden in the Tsukuba Experimental Forest, University of Tsukuba Mountain Science Center (36.11°N, 140.10°E). The trees had originated from 14 wild populations (Supplementary Fig. S1 and Table S1) and were grown from cuttings in the common gardens. Leaves were collected from 3–10 individuals per population (*n* = 108: Tsukuba = 50, Kawatabi = 58). Using scissors surface sterilized with 99% ethanol, leaves (ca. 5 g per tree) were collected in plastic bags, placed in cool bags with ice packs, brought back to the laboratory within 1 day of collection, and stored at 4°C for one to two days.

### Extraction of microbial DNA and amplicon sequencing

DNA of phyllosphere microbes was extracted using the protocol of Tian et al.^82^ with modifications. Collected leaves were soaked in 40 mL of buffer (0.1 M potassium phosphate buffer with 0.1% Tween-80, pH 7.0) and microbes were separated from the leaf surface by sonication for 10 min in a Bransonic ultrasonic bath (Branson Ultrasonics, Danbury, CT, USA). Microbes were precipitated by centrifugation (10,000 ×*g*, 15 min) in a high-speed refrigerated microcentrifuge (Tomy, Nerima, Tokyo, Japan), suspended in 100 μL of PBS and stored at −20°C. DNA was extracted with a Fast DNA Spin Kit for Soil (MP Biomedicals, Irvine, CA, USA) following the manufacturer’s protocol.

Bacterial and fungal DNA was amplified using Ex Taq Hot Start Version (Takara Bio, Kusatsu, Shiga, Japan). The V4 region of bacterial 16S rRNA gene was amplified with the 515F (5′-GTGYCAGCMGCCGCGGTAA-3′) and 806R (5′- GGACTACNVGGGTWTCTAAT-3′) primer set^83^. The PCR mixture (20 μL) contained 1× Ex Taq buffer, 0.2 mM dNTP mixture, 0.05 U Ex Taq HS (Takara Bio, Kusatsu, Shiga, Japan), 0.5 μM forward and reverse primers, 1 μM chloroplast and mitochondria peptide– nucleic acid clumps (to minimize the amplification of chloroplast and mitochondrial 16S rRNA genes from host plants^84^), and 1 μL of the template. PCR was performed as follows: 45 s at 95°C, 28 cycles of 15 s at 95°C, 10 s at 78°C, 30 s at 50°C, 30 s at 72°C, and 5 min at 72°C. The ITS2 region of fungal DNA was amplified with the fITS9 (5′- GAACACAGCGAAATGTGA-3′) and ITS4ngsUni (5′-CCTSCSCTTANTDATATGC-3′) primer set^85,86^. The PCR mixture was as described above except that peptide–nucleic acid clumps were omitted. PCR was performed as above except that the cycles at 78°C were omitted. The second PCR was then performed to add adapter sequences to amplicons. The PCR mixture (20 μL) contained 1× Ex Taq buffer, 0.2 mM dNTP Mixture, 0.05 U Ex Taq HS, 0.5 μM forward and reverse adapters, and 14 μL amplicons from the first PCR. PCR was performed as follows: 45 s at 95°C, 8 cycles of 15 s at 95°C, 30 s at 50°C, 30 s at 72°C, and 5 min at 72°C. Amplicons were purified using NucleoMag NGS Clean-up and Size Select magnetic beads (Macherey-Nagel, Dueren, Germany) following the manufacturer’s protocol. DNA concentration of each sample was measured using Qubit dsDNA Quantification Assay Kits (Thermo Fisher Scientific, Waltham, MA, USA), and an approximately equimolar pool of amplicons was constructed. This pool was commercially sequenced using the Illumina MiSeq platform with the MiSeq Reagent Kit v3 for 2 × 300 bp paired-end reads in a single run (Macrogen, Tokyo, Japan).

### Amplicon sequence processing, ASV classification, and taxonomic assignment

Amplicon sequencing yielded 15,561,772 total raw reads. Sequencing data were analyzed in QIIME 2 v2023.9^87^. Primer sequences of the V4 regions of the 16S rRNA gene and ITS2 region were removed from the 5′ ends of both forward and reverse reads, resulting in a total of 8,467,176 bacterial and 6,799,366 fungal trimmed reads. Quality filtering, chimera removal, and inference of ASVs were performed using the DADA2 pipeline^88^. Bacterial ASVs were inferred from paired-end sequences and fungal ASVs were inferred from single-end sequences, yielding 7,073,527 bacterial and 6,252,014 fungal denoised reads. Taxonomy was assigned to each bacterial ASV using a naïve Bayes classifier^89,90^ based on the SILVA database^91^. The ASVs assigned to chloroplasts or mitochondria and ASVs for which no taxa were assigned at the kingdom level were excluded. Taxonomy was assigned to each fungal ASV using BLAST+^92^ based on the UNITE database^93^. The ASVs assigned to taxa other than fungi at the kingdom level and ASVs that were not assigned to any taxa at the kingdom level were excluded. A total of 3,465,682 filtered bacterial reads (32,090 ± 13,829 reads per sample, mean ± SD) and 5,547,104 filtered fungal reads (51,362 ± 12946 reads per sample, mean ± SD) were used in subsequent analyses.

### Collection of terpenes emitted by Japanese cedar

We collected BVOCs emitted by cedar trees grown in the common garden at Kawatabi Field Center in June 2023; the trees originated from three populations: Ajigasawa (*n* = 9), Azouji (*n* = 10), and Yakushima (*n* = 6). The dynamic branch enclosure system^94^ was used for BVOC collection according to Hiura et al.^50^, except that ambient air was used as the purge gas instead of compressed air from gas cylinders. Briefly, intact branches were enclosed in a Teflon bag with a purge air inlet that supplied ambient air, from which volatile organic compounds and ozone had been removed by charcoal filters. Two air samples (2 L each) per enclosure were collected simultaneously in adsorbent tubes: one for MT measurements and the other one for SQTs and DTs. We followed Hiura et al.^50^ for the analysis of the collected terpenes. Briefly, MT samples were analyzed using a custom-built thermal desorption unit coupled to a gas chromatograph (GC) equipped with a mass selective detector (MSD) and a flame ionization detector (FID) (Agilent 6890/5973, Agilent, Santa Clara, US). SQT and DT samples were analyzed using a GC/MSD (Agilent 6890/5973, Agilent, Santa Clara, US) following solvent extraction and concentration.

To standardize emission rates, we applied the G93 model^95^ and converted the measured emission rates to basal emission rates, which were defined as the emission rates at a standard leaf temperature of 30 °C, using the values of the constant *β* for models of the rates of terpene emission by *C. japonica* reported previously^48,96^. For statistical analysis, the basal emission rates were log-transformed after adding 0.01 (<1/100th of the lowest measured basal emission rate) to avoid zero or negative values. Data of emitted terpenes are shown in Supplementary Table S3.

### Acquisition and analysis of climatic data at the original locations of the wild Japanese cedar populations

The following climate data at the original locations of the wild populations of *C. japonica* for 1981–2010 were obtained from the CHELSA database^59–62^: annual mean temperature (bio1), standard deviation of the monthly mean temperatures (bio4), annual precipitation (bio12), coefficient of variation of the monthly precipitation estimates (bio15), mean monthly near-surface relative humidity (hurs_mean), mean monthly potential evaporation (pet_penman_mean), monthly mean temperatures (tas01–12), and monthly precipitation (pr01–12). The sum of the monthly precipitation for the months with mean temperatures below 0 °C was calculated for each location and used as an index of snowfall (snow). This index and the six average climatic variables other than the monthly means were summarized into PCs by PCA using the prcomp function in R v4.3.3^97^.

### Statistical analysis

Data analysis and visualization were performed using the tidyverse^98^, phyloseq^99^, vegan^100^, MASS^101^, and ggplot2^102^ packages in R v4.3.3^97^. Using the vegan^100^ package, the ASV table was rarefied to the minimum reads per sample to ensure equal sequencing depth (bacteria: 9,099 reads, fungi: 17,427 reads). The ASV richness of bacteria and fungi was calculated using the specnumber function in the vegan^100^ package. Bray–Curtis dissimilarities among the microbial communities were calculated to measure their beta diversity using the vegan^100^ package. Nonmetric multidimensional scaling (NMDS) based on Bray–Curtis dissimilarities was performed to visualize beta diversity. A permutational multivariate analysis of variance (PERMANOVA) with the Bray–Curtis dissimilarities as the response variable was performed to test for differences in microbial community composition among common gardens and populations, which were included together with their interactions as explanatory variables in the test. PERMANOVA was also performed using the climatic PC1 and PC2 at the origin sites of host individuals as explanatory variables instead of their populations to test for correlation between microbial community composition and climate at the origin sites of host individuals. The list of fungal pathogens of Japanese cedar throughout Japan was obtained from Kobayashi^63^. Fungal ASVs of the same genus as pathogens in the list were selected as putative pathogenic fungi. To explore the drivers of the total relative abundances of the putative pathogenic fungi, a negative binomial generalized linear model was fitted with the aggregated abundances of these fungi after rarefaction as the response variable using the glm.nb function with a log link function in the MASS^101^ package. The climatic PC1 and PC2 at the origin sites of host individuals and common garden locations were included in the model as explanatory variables. The significance of the explanatory variables was tested by the Wald test.

To analyze the terpene emission profiles of Japanese cedar, we performed PCA of the basal emission rates of terpenes after log transformation using the prcomp function. To investigate the relationship between the terpene emission profile and the local climate at the origin sites of the cedar trees, we fitted linear regression models with the terpene PCs as the response variable. Simple linear regression models were fitted with either the terpene PC1 or PC2 as the response variable and either the climatic PC1 or PC2 at the origin as the explanatory variable, resulting in four models. The significance of the explanatory variables in each model was tested by Student’s *t*-test.

The following procedure was used to analyze the relationship between the basal terpene emission rate and phyllosphere microbial community composition. First, to avoid a large distortion of the entire NMDS plot due to the microbial community outliers, we detected them using CLOUD analysis^103^, a non-parametric test that is based on a distance matrix between microbial communities and detects microbial samples whose mean distance to neighbors is significantly different from those of other samples. Outliers in the bacterial and fungal communities were detected using the mean Bray–Curtis dissimilarity to five neighbors. The only detected outlier (the fungal community of one tree from the Azouji population) was excluded from the subsequent analysis. Next, NMDS based on the Bray– Curtis dissimilarities was performed only for those individuals from which emitted terpenes were collected. Correlation was then tested between the NMDS coordinates constructed from the phyllosphere microbial community compositions and the basal emission rates of terpenes by vector fitting in the vegan package^100^. The basal emission rate of each terpene, their total sum, or the sum for each category (i.e., MT or SQT) was used for the vector fitting after log transformation. The sum for DT was not used for the analysis since only one tree emitted it. Correlation was also tested between the abundances of bacterial phyla or fungal classes and the NMDS coordinates constructed from the community compositions by vector fitting.

## Data availability

The datasets generated and analysed during the current study are available in the DDBJ Sequence Read Archive under the accession number PRJDB20421. All R scripts used for the data analysis are available on GitHub (https://github.com/Satoyoshi-ISHIZAKI/CedarPhyllosphere_SciRepo.git).

## Acknowledgements

This research was partly supported by the Japan Society for the Promotion of Science to T.I.K. (No. 22H05715), Y.S. (No. 23H04970), and T.H. (Nos. 21H02227, 21H05316, 24K01809). We thank the staff of Kawatabi Field Science Center, Tohoku University and Mountain Science Center, University of Tsukuba for supporting our fieldwork.

## Author contributions

S.I. performed statistical analysis and wrote the first draft; T.I.K and T.H. generated hypotheses and designed research; S.I. and T.I.K. collected and performed experiments with microbial samples; Y.O. sampled terpenes emitted from the Japanese cedars; Y.O. and T.S. analyzed the collected terpenes; Y.S. and Y.T. designed and maintained the common gardens. All authors participated in manuscript preparation.

