## Supplementary Information for "Common garden experiments suggest terpene-mediated interactions between phyllosphere microbes and *Cryptomeria japonica*"

**Terpene emission mediates interactions between phyllosphere microbes and host population origins in *Cryptomeria japonica***

Satoyoshi Ishizaki^1,^*, Tetsuo I. Kohyama^1^, Yuki Ota^1^, Takuya Saito^2^, Yoshihisa Suyama^3^, Yoshihiko Tsumura^4^, Tsutom Hiura^1^

^1^ Department of Ecosystem Studies, Graduate School of Agricultural and Life Sciences, The University of Tokyo, Tokyo 113‑8657, Japan

^2^ National Institute for Environmental Studies, Tsukuba 305‑8506, Japan

^3^ Field Science Center, Graduate School of Agricultural Science, Tohoku University, Osaki 989‑6711, Japan

^4^ Institute of Life and Environmental Sciences, University of Tsukuba, Tsukuba 305‑8572, Japan

* Corresponding author: Satoyoshi Ishizaki

This file includes: Figures S1 to S3; Tables S1 to S3; SI references

Supplementary Figures

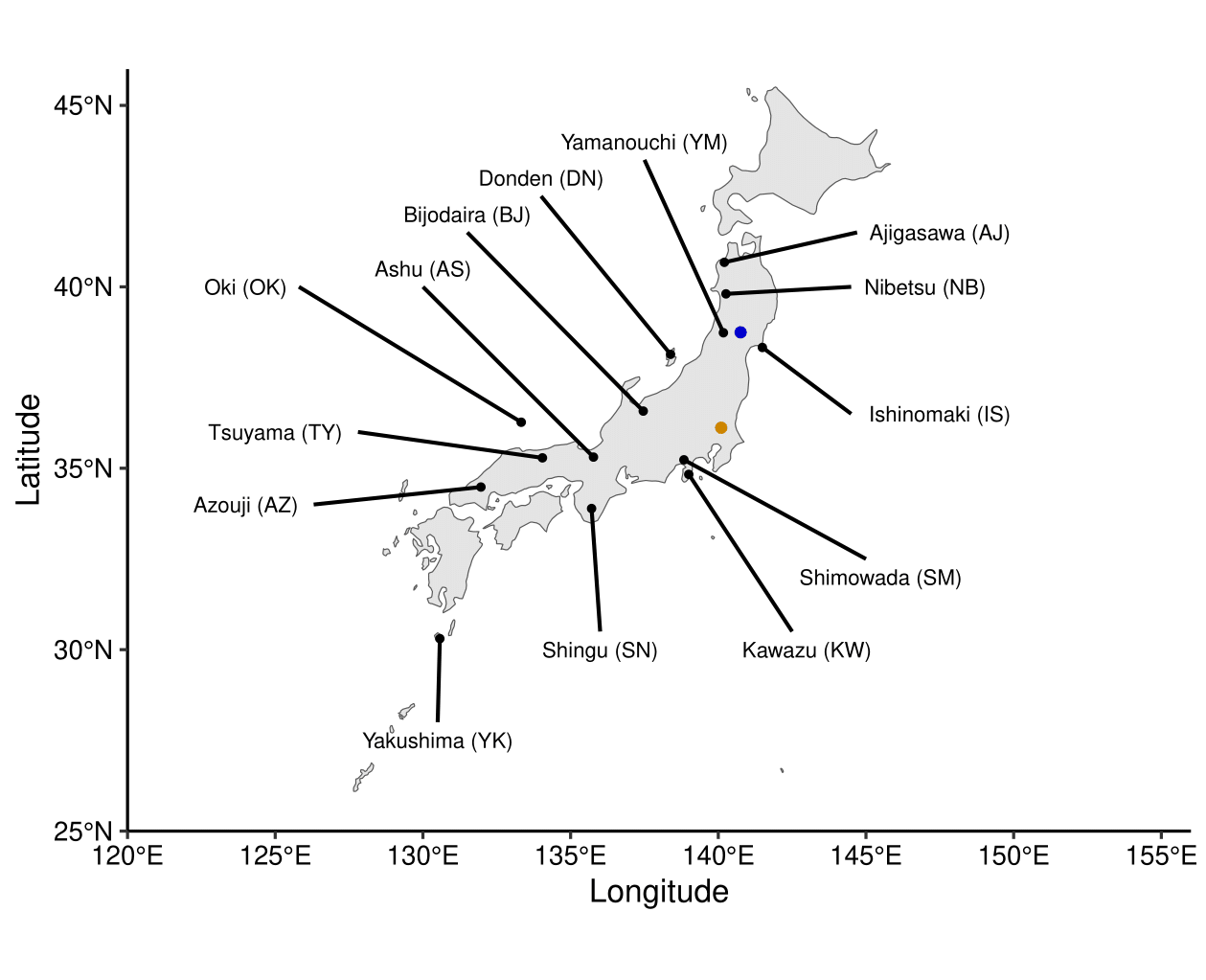

Figure S1.

Locations of the common gardens (blue: Kawatabi, orange: Tsukuba) and the original wild populations of *Cryptomeria japonica* planted in the common gardens. The map was generated using ggplot2^1^, sf^2^, rnaturalearth^3^, and rnaturalearthdata^4^ packages in R 4.3.3^5^ using Natural Earth 4.1.0 (<https://www.naturalearthdata.com>).

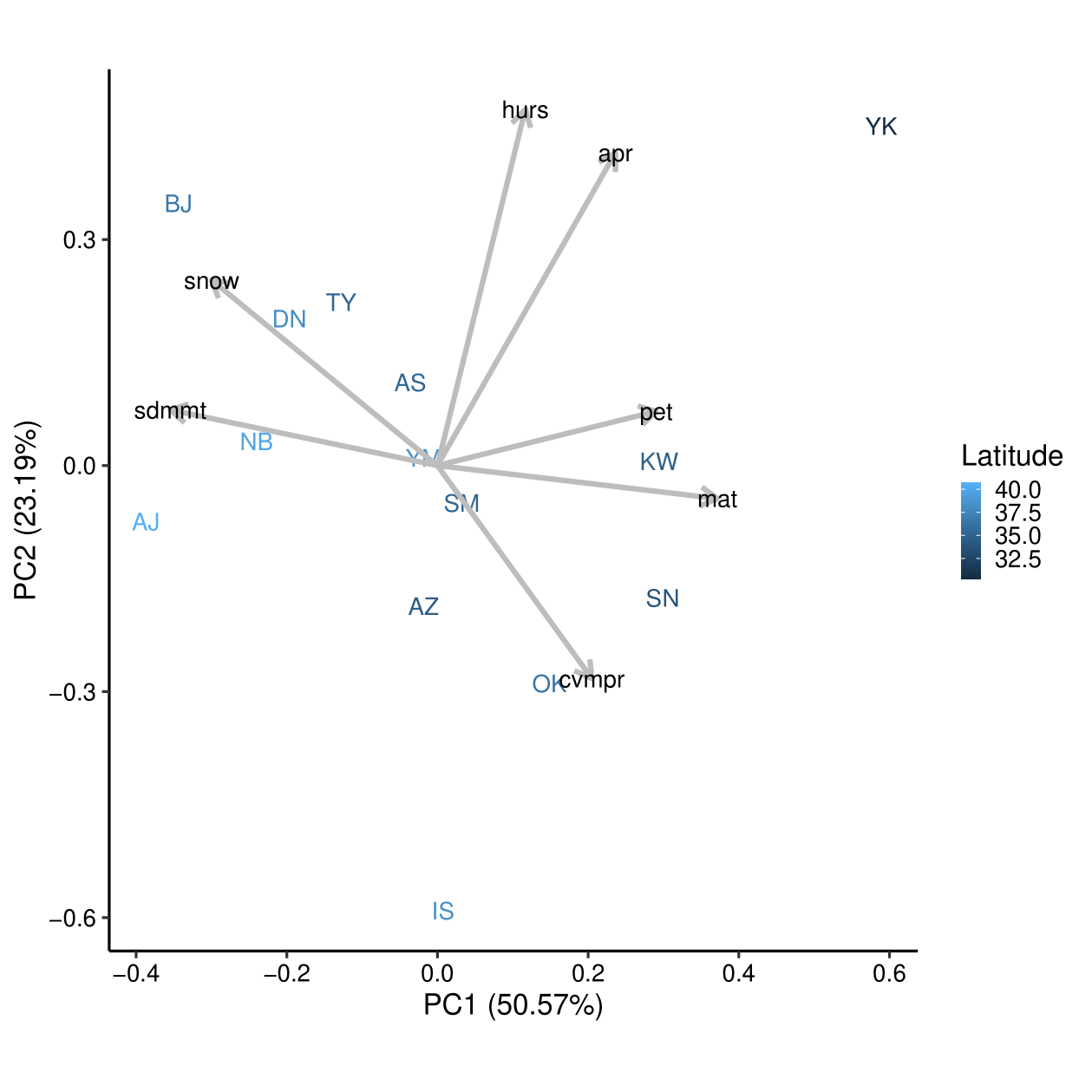

Figure S2.

Principal component analysis of seven climate factors at the original locations of the wild populations of *Cryptomeria japonica*. Mean annual temperature (mat), standard deviation of the monthly mean temperatures (sdmmt), annual precipitation (apr), coefficient of variation of the monthly precipitation estimates (cvmpr), mean monthly near-surface relative humidity (hurs), mean monthly potential evaporation (pet), and the sum of the monthly precipitation for the months with mean temperatures below 0 °C (snow) were used for the analysis. Climatic data for 1981–2010 were obtained from the CHELSA database^6–9^.

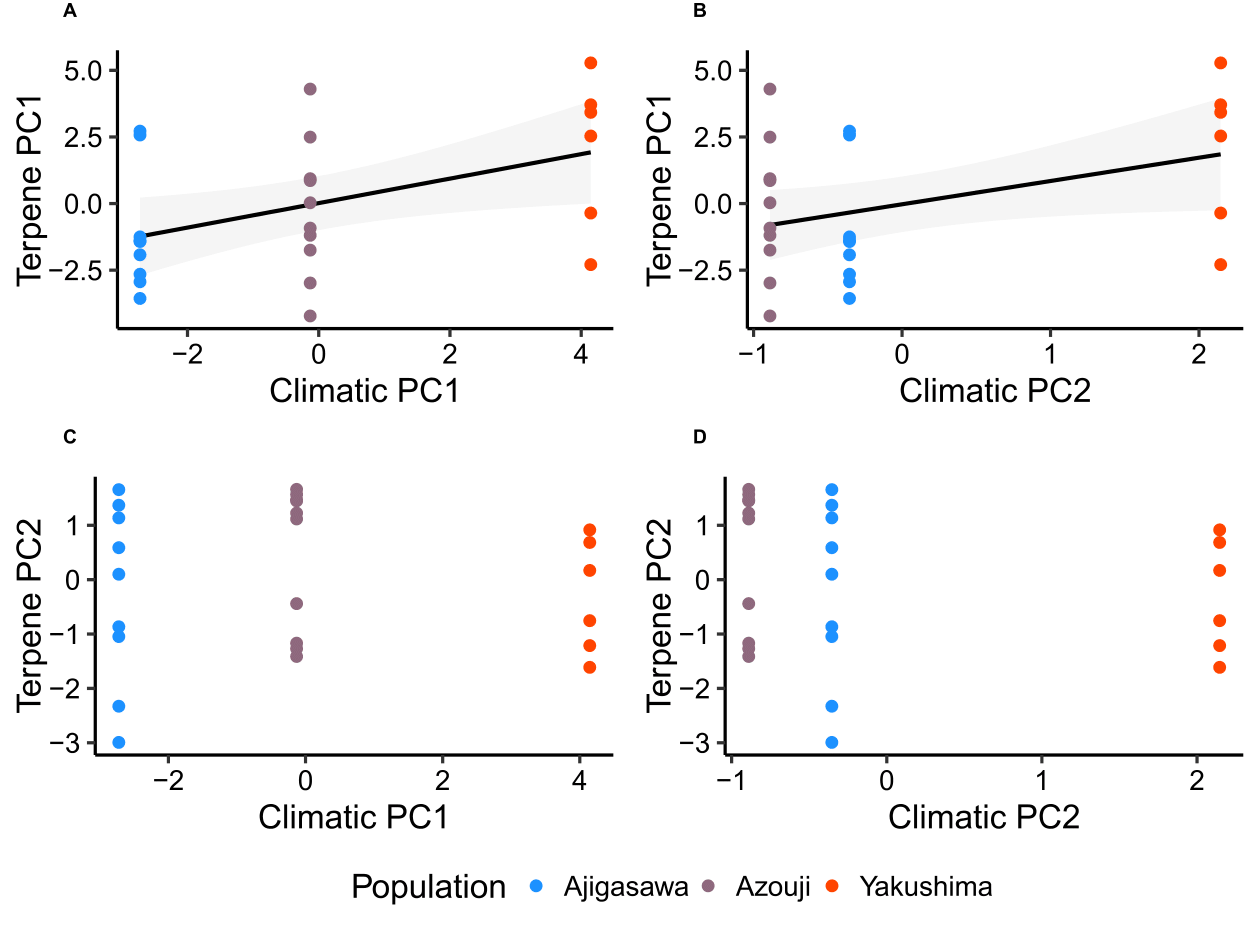

Figure S3.

Relationships between the terpene PCs and the climatic PCs at the origin sites of the cedar individuals. Colors indicate the populations of the cedars. Solid lines represent fitted values from linear models where the terpene PCs were modeled as a function of the climatic PCs. Shaded areas indicate the 95% confidence intervals of the model predictions.

Table S1.

Locations of the 14 wild populations of *Cryptomeria japonica*

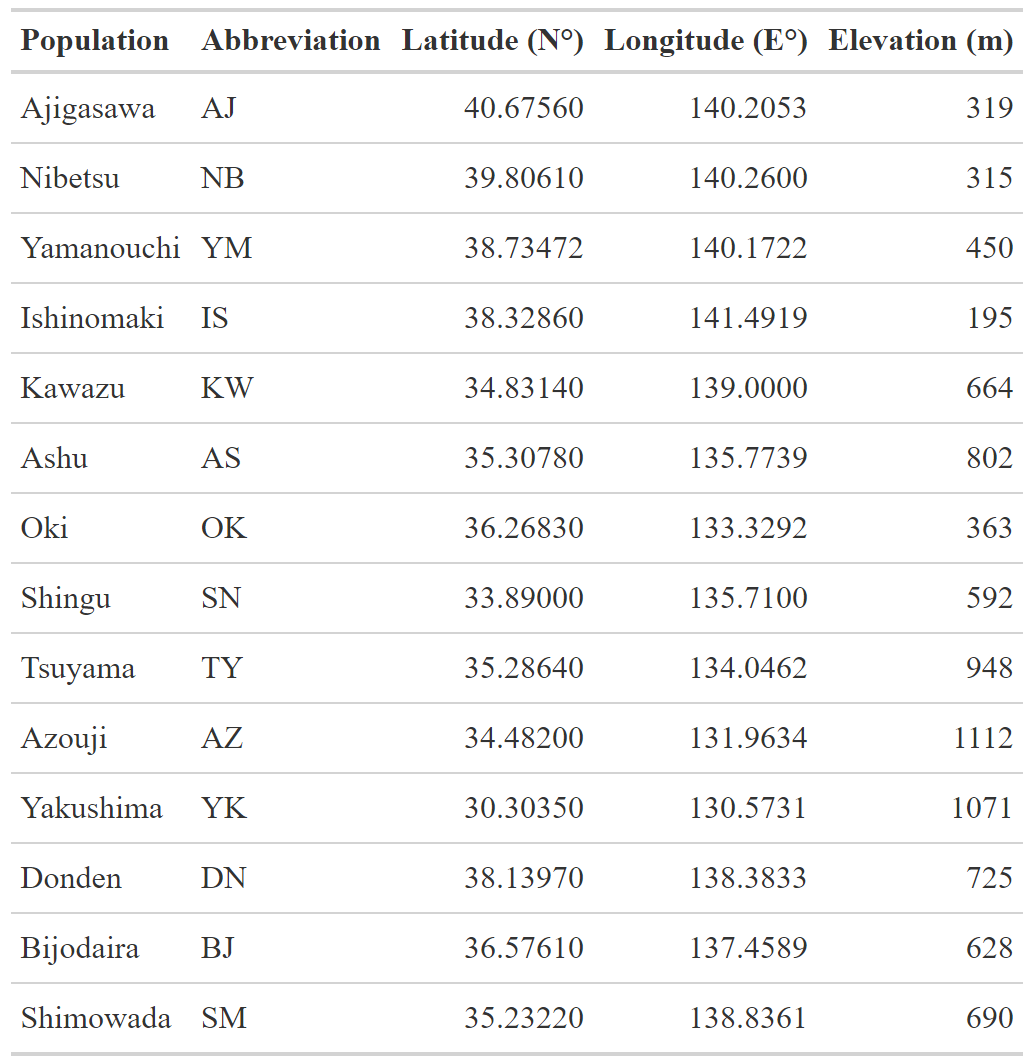

Table S2.

Putative pathogenic fungal genera detected in the amplicon sequencing data. Some species in these genera have been described as cedar pathogens^10^

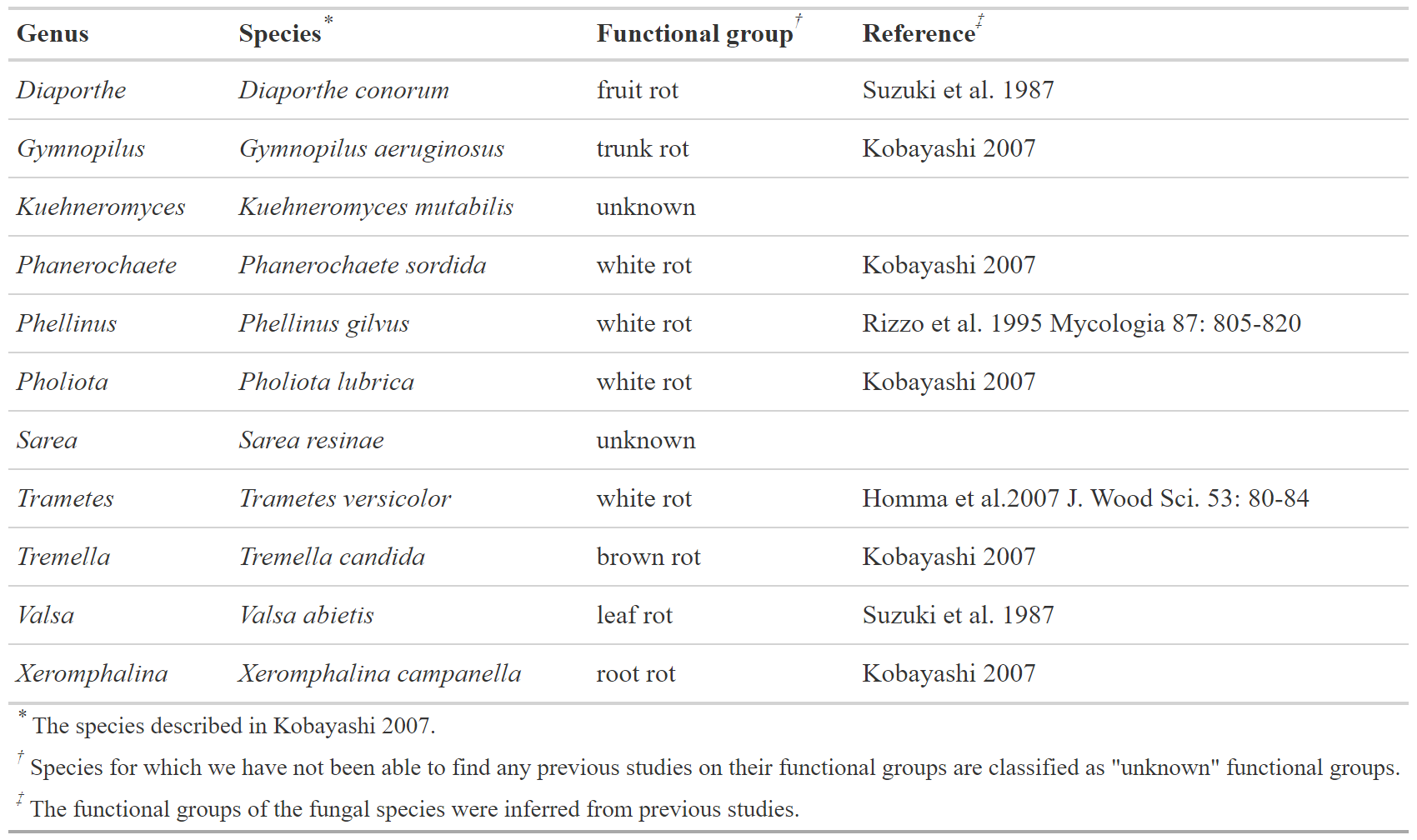

Table S3.

Basal emission rates of terpenes (ng gdw^−1^ h^−1^) in *Cryptomeria japonica* grown in the common gardens. HN, LD, and ST are subpopulations in the Yakushima population.

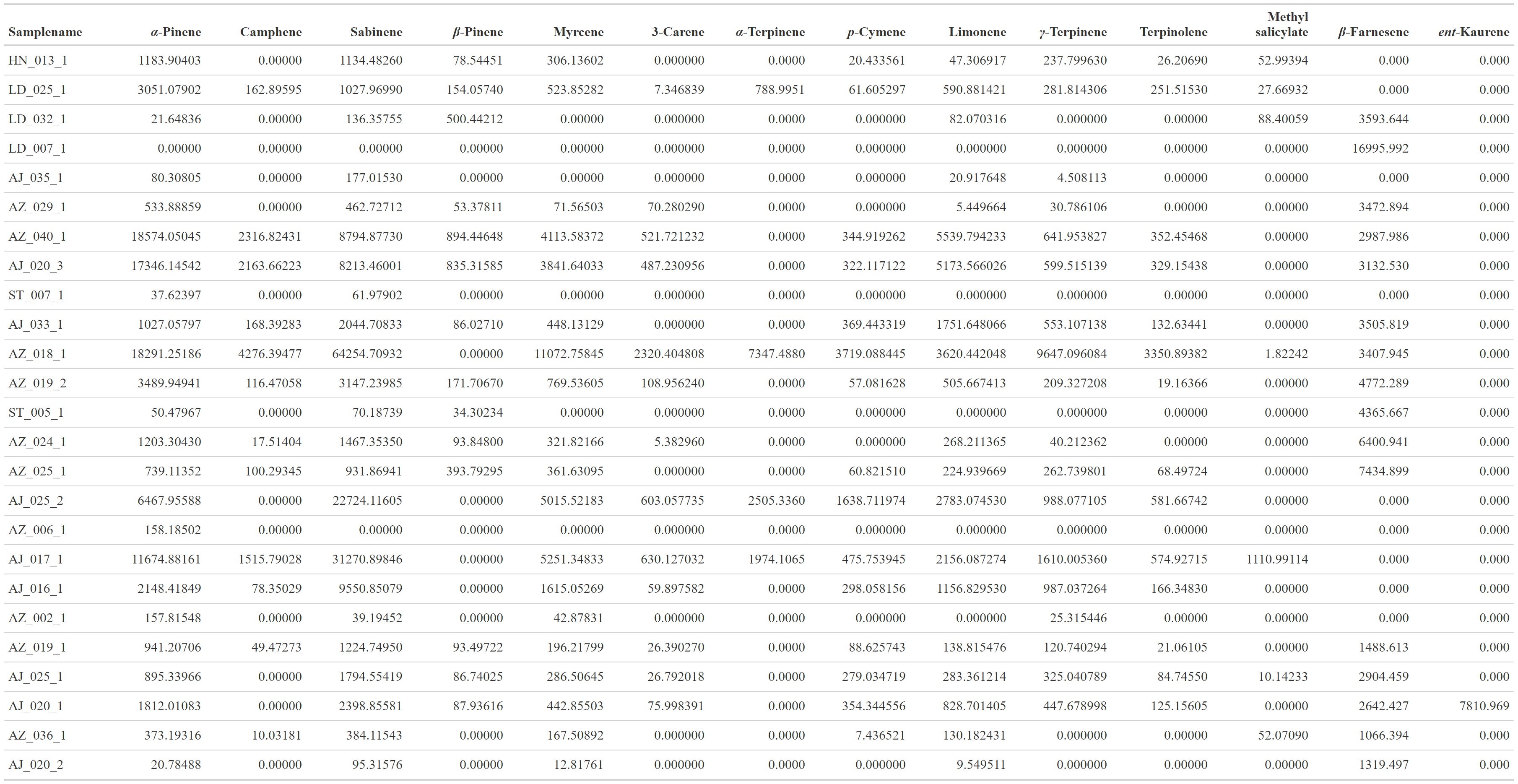
